# Breaking the “one nucleus, one whole genome” rule: *Neurospora crassa* separates its haploid chromosomes into different nuclei

**DOI:** 10.1101/2025.11.08.687325

**Authors:** Jinyi Tan, Yan Xu, Piao Yang, Shuai Huang, Monika Fischer, Yuelin Zhang, Xin Li

## Abstract

The nucleus is a defining feature of eukaryotic cells, compartmentalizing the genome to enable precise regulation of diverse cellular processes. The long-standing paradigm in textbooks states that each nucleus carries at least a complete haploid genome —the “one nucleus, one whole genome” rule. A striking exception was recently uncovered: in two sclerotia-forming plant pathogenic fungi in the Sclerotiniaceae family, *Sclerotinia sclerotiorum* and *Botrytis cinerea*, haploid chromosomes are irregularly distributed across multiple nuclei. However, whether such phenomenon occurs in other eukaryotes, particularly in non-pathogenic fungi, is unclear. Through a literature survey, we identified many fungi that can potentially also separate their haploid chromosomes across different nuclei. Interestingly, *Neurospora crassa*, a foundational fungal genetics model, came out as a candidate. Its conidia, typically containing two or three nuclei, were believed to harbor a complete haploid genome within each nucleus. However, using approaches including chromosome counting, flow cytometry, and fluorescence in situ hybridization, we found that haploid chromosomes in *N. crassa* can be unevenly distributed among multiple nuclei. These findings challenge longstanding assumptions in fungal genetics and provide new directions for investigating genome organization. Our results indicate that the organisms violating the “one nucleus, one whole genome” rule are beyond sclerotia-forming pathogenic fungi. Since over 90% of fungi have not even been described, we propose that many more of them may likewise partition their genomes non-uniformly across nuclei, a process that could facilitate their adaptation and evolution.

## Introduction

A defining feature of eukaryotes is the presence of nuclei, which compartmentalize genetic material and orchestrate gene expression. It has been long held that each nucleus contains at least one full set of chromosomes required for proper genomic function and cellular development – an assumption referred to as the “one nucleus, one whole genome” rule. However, our recent study challenges this long-standing principle. Using chromosomal counting, fluorescence in situ hybridization (FISH), flow cytometry-based DNA quantification, and single-nucleus polymerase chain reaction (PCR) analyses, we uncovered that in two plant pathogenic fungi belonging to the Sclerotiniaceae family –*Sclerotinia sclerotiorum* and *Botrytis cinerea* – haploid chromosomes are distributed among different nuclei (1).

Such finding raises the next question of how widespread this unconventional mode of chromosomal organization is. Both *S. sclerotiorum* and *B. cinerea* produce bi-or multinucleate spores. Interestingly, they also generate uninucleate spermatia that are not viable, likely due to the incomplete chromosome sets within (2,3). To explore whether similar chromosomal partitioning pattern might be more widespread, we undertook an extensive literature survey focusing on fungal species reported to produce bi-or multinucleate spores along with a high frequency of nonviable uninucleate cells (4). These species were considered likely candidates that may also be breaking the “one nucleus, one whole genome” rule. Interestingly, *Neurospora crassa* came out as a prime candidate. Although it can produce viable uninuclear microconidia, more than 60% are nonviable, possibly due to the absence of a complete chromosome complement (5,6).

*N. crassa* is a non-pathogenic ascomycete that has long served as a classical model organism in fungal genetics and molecular biology, contributing to landmark discoveries such as the “one gene, one enzyme” hypothesis (7,8). Owing to its well-characterized genetics and ease of manipulation, *N. crassa* was almost the first filamentous fungus to have its genome fully sequenced, establishing it as a preferred model for genome-wide functional studies and for numerous discoveries related to chromosomal behavior and gene regulation (9–11). Its genome, approximately 40 Mb in size, is organized into seven linkage groups as determined through pulsed-field gel electrophoresis (PFGE) and whole-genome sequencing and assembly (9,10,12). In this short report, through chromosome counting, flow cytometry and fluorescence in situ hybridization (FISH) analyses, we show that *N. crassa* can partition its haploid genome across multiple nuclei, thereby extending the scope of breaking the “one nucleus, one whole genome” rule beyond the Sclerotiniaceae family.

## Results

*N. crassa* produces abundant asexual spores, known as conidia, which are typically multinucleate. When we grew *N. crassa* on potato dextrose agar (PDA), majority of fresh conidia contain 2 to 3 nuclei (Figure 1A), whereas conidia from older cultures show an obvious increase in nuclear number, with a higher proportion containing more than three nuclei (Figure 1B and C). This indicates that genomes within the conidia are not predefined; rather, they can undergo endoreduplication through time and become more variable. To quantify the DNA amount in each nucleus, we employed flow cytometry through fluorescence-activated cell sorting (FACS) using purified nuclei from fresh mycelia. Haploid and diploid *Saccharomyces cerevisiae* strains BY4741 and BY4743, and the filamentous plant pathogen *S. sclerotiorum* were used as controls. Based on the relative fluorescence intensity (RFI) of nuclei (Figure 2A and B), each nucleus contained about a quarter of its full haploid genome, suggesting that each nucleus within *N. crassa* mycelia harbors only parts of its full genome (Figure 2C).

**Figure 1.**
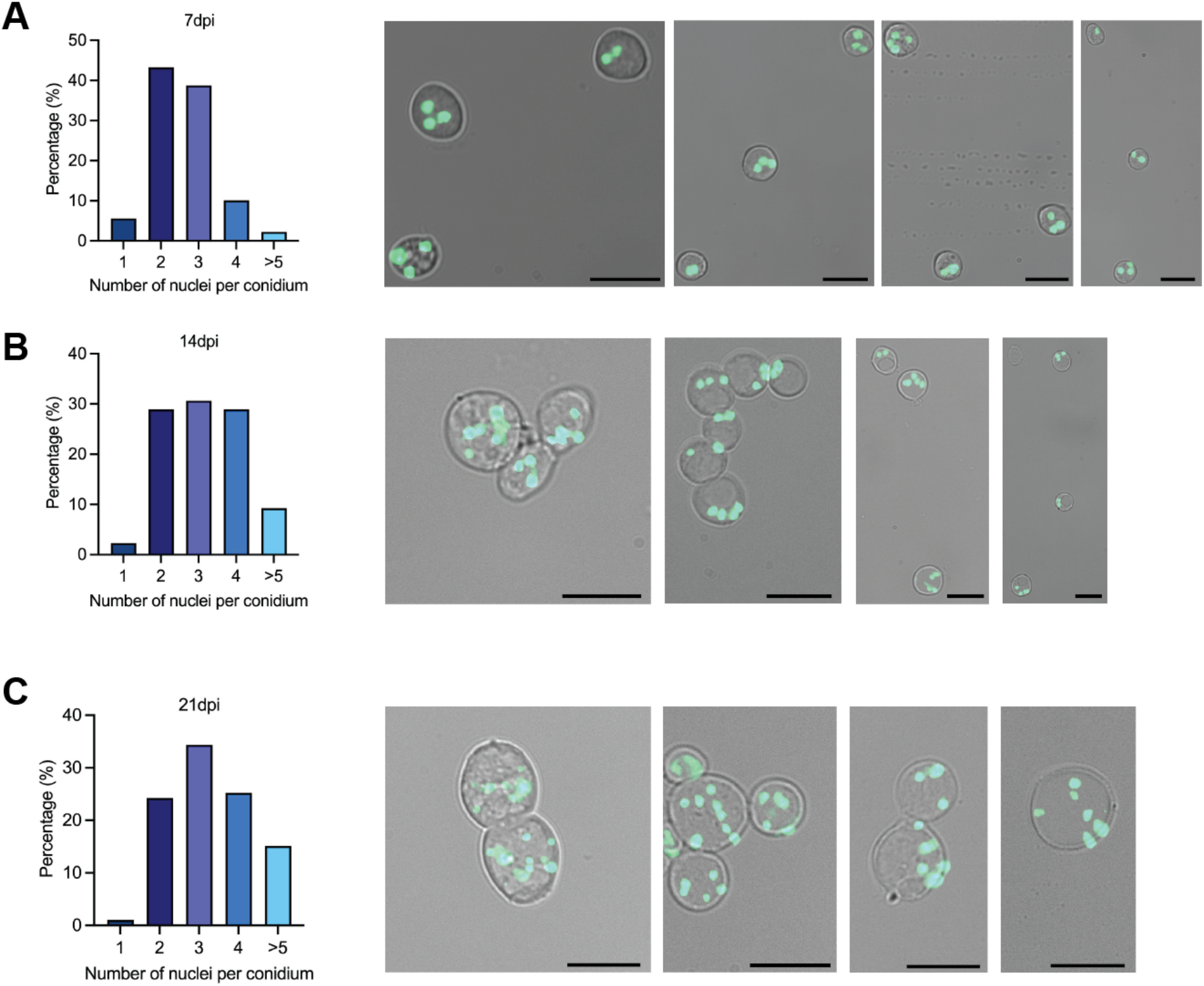
*N. crassa* conidia have multiple nuclei, with nuclear number increasing as culture ages. **(A)** Quantification of conidial nuclei number in fresh culture at 7 days post-inoculation on PDA (7 dpi) (left). Representative fluorescence microscopy images of conidial nuclei in the hH1-sGFP strain are shown (right). Scale bar, 10 μm. **(B)** Quantification of conidial nuclei number under the same condition at 14 dpi (left). Representative fluorescence microscopy images of conidial nuclei in the hH1-sGFP strain are shown (right). Scale bar, 10 μm. **(C)** Quantification of conidial nuclei number under the same condition at 21 dpi (left). Representative fluorescence microscopy images of conidial nuclei in the hH1-sGFP strain are shown (right). Scale bar, 10 μm.

**Figure 2.**
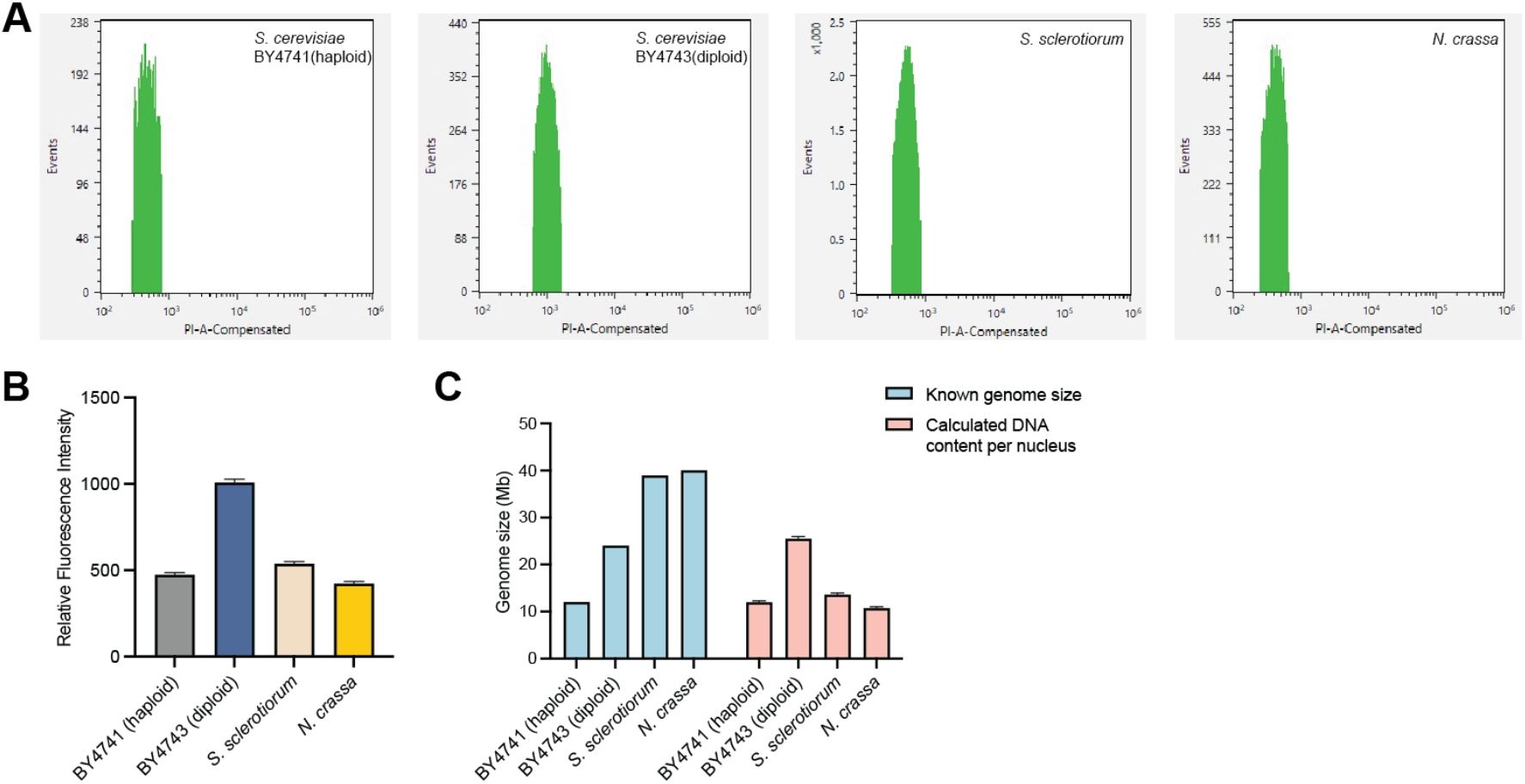
On average, each nucleus of *N. crassa* contains a portion of its haploid genome. **(A)** Nuclei sorting by FACS. The isolated nuclei from *Saccharomyces cerevisiae* BY4741 (haploid), BY4743 (diploid), *S. sclerotiorum* and *N. crassa* mycelia were flowed into the Sony MA900 cell sorter. The selected events were plotted with the number of events against the PI fluorescence intensity. **(B)** Propidium iodide (PI) fluorescence intensity of sorted purified nuclei from *S. cerevisiae* BY4741, BY4743, *S. sclerotiorum* and *N. crassa* mycelia. The PI fluorescence intensity values were calculated based on the mean fluorescence intensity of each individual well. Error bars indicate the standard deviation of the values from 15 wells. **(C)** Calculated DNA content per nucleus in *S. cerevisiae* BY4741, BY4743, *S. sclerotiorum* and *N. crassa*. The known genome sizes of these strains are 12, 24, 38.9 and 40 Mb, respectively. The DNA content in each nucleus was calculated by multiplying the genome size of the haploid yeast strain BY4741 by the fold-change values derived from Fig 2B. Error bars indicate the standard deviation (*n* = 15).

We then counted chromosomes inside the protoplasts isolated directly from the conidia. As observed in *S. sclerotiorum*, where most ascospores harbor approximately 16 chromosomes (1), the largest percentage of *N. crassa* conidia contains around seven chromosomes (Figure 3A, B and E). However, substantial proportions harbor more than seven chromosomes, with counts reaching up to 22. These observations suggest that aneuploidy may be more common in *N. crassa* conidia, and that conidia containing equal to or more than 14 chromosomes could have arisen through endoreduplication (Figure 3D and E). Live-cell observation of the *N. crassa* strain carrying the hH1-sGFP (green fluorescence protein fused with histone H1) transgene with confocal fluorescence microscopy further confirmed substantial variation in chromosome number among nuclei. Although precise chromosome counts are difficult to determine using hH1-sGFP fusion strains, the observed numbers were largely fewer than seven (see S1 Movie, showing division of a nucleus with about two chromosomes, and S2 Movie, showing division of a nucleus with approximately four chromosomes). Together, these data support that most viable conidia are haploid and can separate their haploid genome into different nuclei.

**Figure 3.**
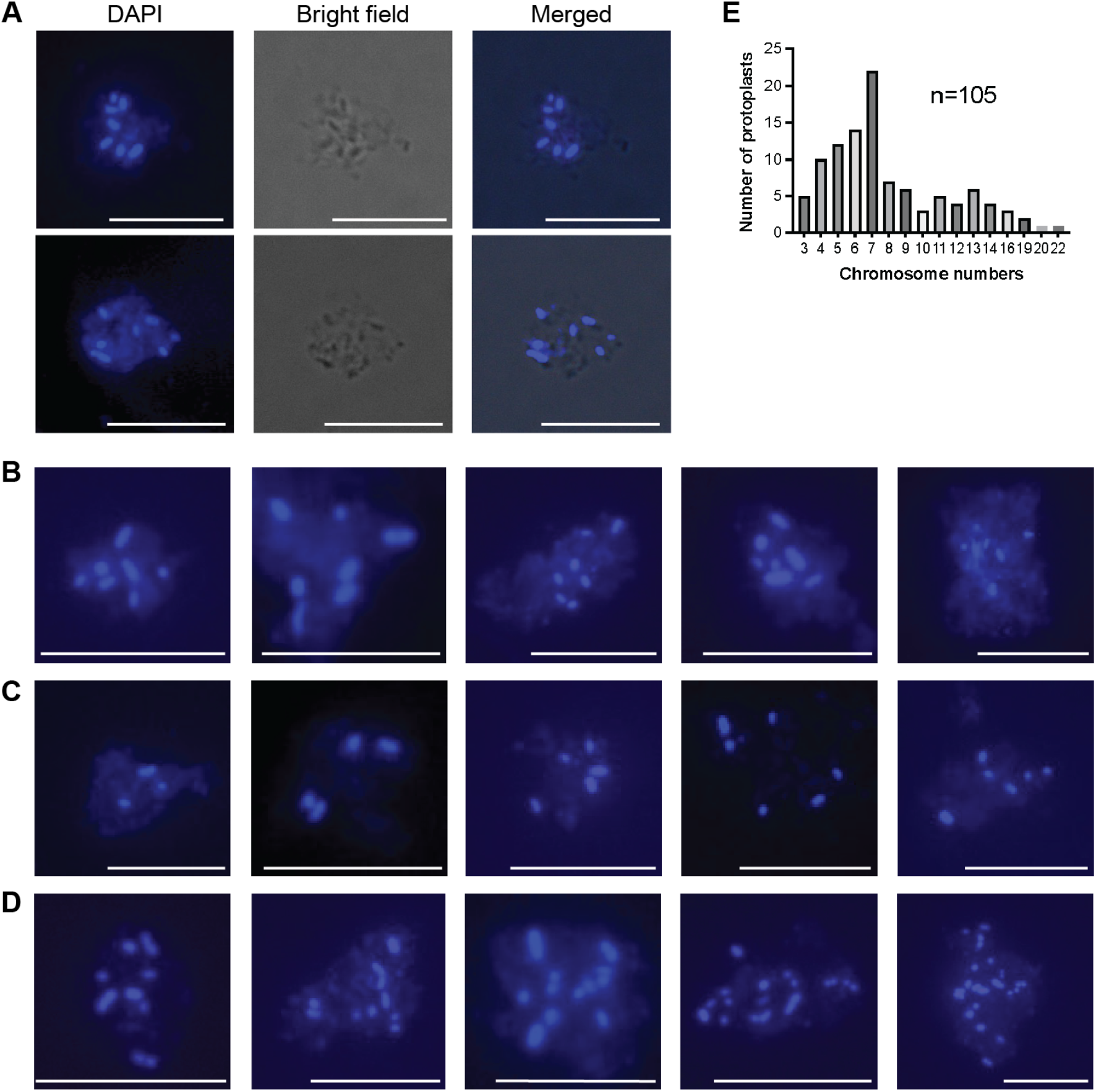
Chromosome counts from *N. crassa* conidial protoplasts. **(A)** DAPI stained chromosomes inside two representative conidial protoplasts. Scale bar, 10 μm. **(B)** DAPI-stained conidial protoplasts showing seven chromosomes. Scale bar, 10 μm. **(C)** DAPI-stained conidial protoplasts showing fewer than seven chromosomes. Scale bar, 10 μm. **(D)** DAPI-stained conidial protoplasts showing more than seven chromosomes. Scale bar, 10 μm. **(E)** Distribution histogram of chromosome counts from a total of 105 protoplasts of *N. crassa* conidia.

Lastly, to corroborate our observations, we performed fluorescence in situ hybridization (FISH) using linkage group I (LG I)- and linkage group IV (LG IV)- specific probes (Figure 4A). When these probes were applied individually, hybridization signals were consistently detected in only one nucleus per conidium (Figure 4B and C). Importantly, probe signals were never detected simultaneously in multiple nuclei within the same conidium (Figure 4B, C and D). As a control, a telomere repeat probe that is present in all linkage groups yielded signals mostly in all nuclei (Figure 4A), confirming that probe accessibility was not limiting (Figure 4B, C and D). Interestingly, the intensity of the fluorescence signals from the telomere repeat probe can be drastically different among nuclei within the same cell, likely reflecting the different number of chromosomes within (Figure 4B and C). The intensity of the signals also seems to correlate with the size of the nuclei. A similar trend was observed in live-cell imaging using the hH1-sGFP strain, where nuclei containing fewer chromosomes appeared smaller than those harboring more chromosomes. Thus, haploid chromosomes in *N. crassa* can partition unevenly in these bi-or tri-nuclear conidia.

**Figure 4.**
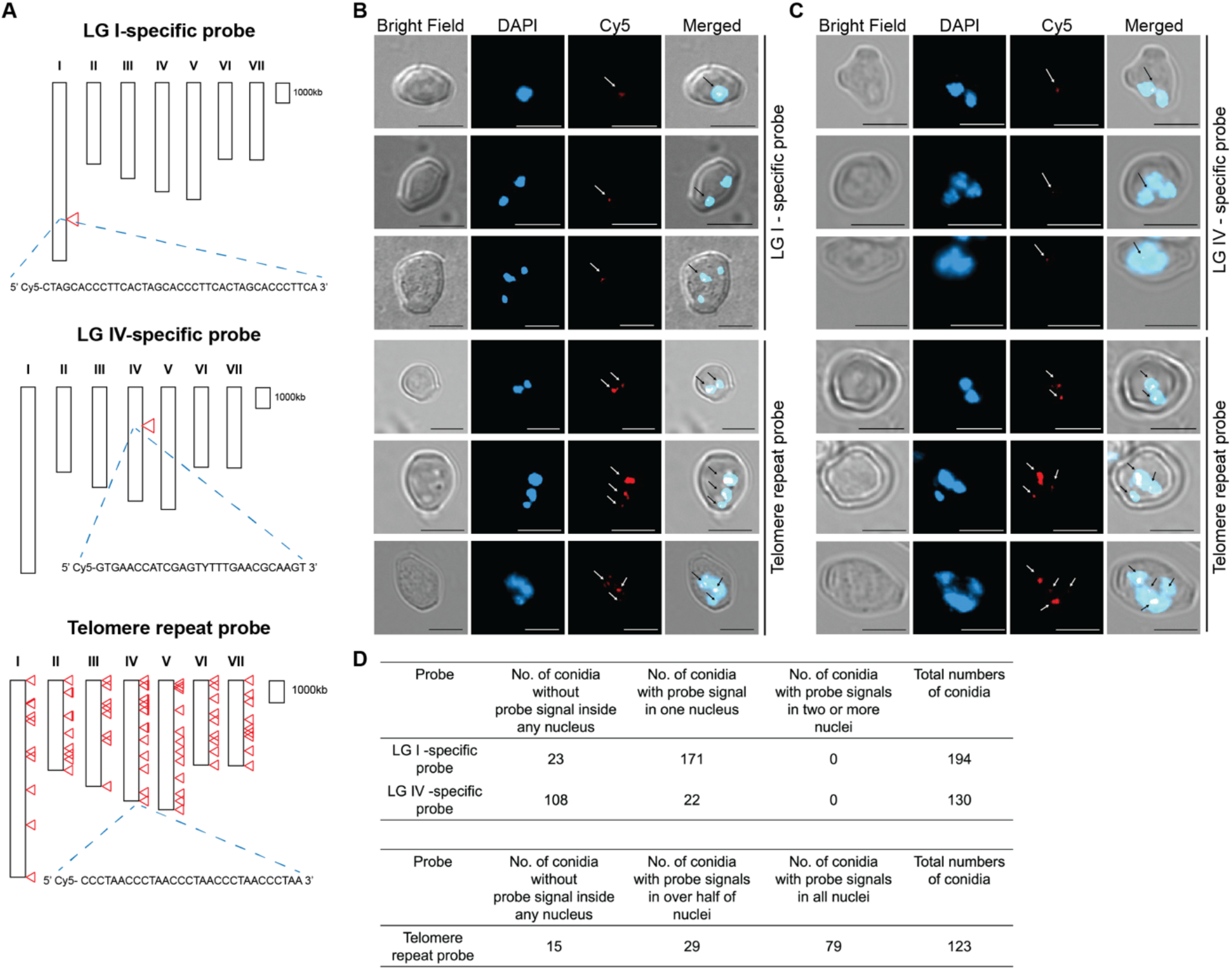
A *N.crassa* conidium can separate its haploid chromosomes into one, two, or three nuclei. **(A)** The hybridization sites of the LG I-specific (top), LG IV-specific (middle) and telomere repeat (bottom) probes on the *N. crassa* chromosomes as indicated by red triangles. **(B)** FISH signals of LG I-specific (top) and non-specific (bottom) probes in representative conidia. Blue and red signals are from DAPI-stained nuclei and hybridized fluorescent probe, respectively. The hybridized Cy5 probe signals are pointed out by arrows. Scale bars, 5 μm. **(C)** FISH signals of LG IV-specific (top) and non-specific (bottom) probes in representative conidia. Blue and red signals are from DAPI-stained nuclei and hybridized fluorescent probe, respectively. The hybridized Cy5 probe signals are pointed out by arrows. Scale bars, 5 μm. **(D)** Summary of FISH results for LG I-, LG IV-specific and telomere repeat probes. Among conidia with a single LG I-specific signal, 6% were uninucleate, 48% contained two nuclei, 35% contained three nuclei, and 11% contained four or more nuclei. For conidia with a single LG IV-specific signal, 22 contained two nuclei and 5 contained three nuclei. The lower number of conidia exhibiting a LG IV-specific probe signal is likely due to reduced probe hybridization efficiency or limited accessibility caused by the presence of a degenerate nucleotide in the probe.

## Discussion

Our data here support the conclusion that *N. crassa* conidia can partition their haploid chromosomes among different nuclei, expanding the range of species in which genome partitioning across nuclei has been observed. This feature will offer a major practical advantage. For gene replacement experiments to knock out or modify target genes, fresh conidia are already being used, skipping the step of uninucleate microconidia purification (13). The same is true for mutagenesis in forward genetic analyses (14). Just as with *S. sclerotiorum*, binucleate ascospores or multinuclear conidia of *N. crassa* can be used with UV or chemical mutagens to generate pure haploid mutants (15).

Interestingly, uninucleate *N. crassa* microconidia exhibit a low germination rate of about 10-30% (6,16), whereas uninucleate spermatia cells in *S. sclerotiorum* and *B. cinerea* are completely nonviable likely due to the incomplete genomes within (2,3). We believe the reason why a small fraction of the uninucleate microconidia is viable in *N. crassa* is probably due to its low haploid chromosome number. Its seven chromosomes may occasionally be packed into just one nucleus, allowing the spore to be viable. As for *S. sclerotiorum* and *B. cinerea*, 16 or 18 chromosomes are seemingly almost impossible to be packed simultaneously into a single nucleus during nuclear division.

The chromosomal and nuclear states in *N. crassa* conidia appear to be more complex than those of haploid *S. sclerotiorum* ascospores. Based on our chromosome counting results, although the largest percentage of *N. crassa* conidia have seven chromosomes, a substantial fraction contains more. Together with the observed increase in number of nuclei inside the conidia through time, these data suggest that some *N. crassa* conidia may be aneuploid, while others may contain two or more chromosome sets as a result of mitotic divisions occurring after initial conidial formation. Similar pattern could exist with multinuclear conidia of other fungi, such as *B. cinerea*. Such genomic flexibility in conidia may help the survival and quick adaptation of the organism.

In summary, our findings in *N. crassa* suggest that dividing haploid chromosomes across multiple nuclei may be more widespread than previously recognized. It is not restricted to sclerotia-forming plant fungal pathogens of the Sclerotiniaceae family. Other Ascomycetes and Basidiomycetes with known bi-or multi-nucleate ascospores and basidiospores, such as *Morchella, Oudemansiella* and *Cortinarius*, also warrant further investigations (17–19). Studying these organisms could provide important insights into the extensiveness of this phenomenon and advance our understanding of chromosome biology and genome evolution. Moreover, the precise molecular mechanisms underlying such chromosome splitting phenomenon remain puzzling and represent an intriguing new area for future research. For instance, how do different nuclei in the same cell communicate and coordinate with each other? What are the key genes and proteins involved in nuclear coordination? Are there special checkpoints to ensure proper chromosomal distribution? Advanced live-cell imaging and single-nuclei sequencing could be particularly valuable for tracking these processes in single cells.

## Materials and Methods

### Fungal strains and culture conditions

*N. crassa* WT strains 74-OR23-1VA (FGSC#: 2489) and ORS-SL6a (FGSC#: 4200), as well as the hH1-sGFP strains (20) were grown on potato dextrose agar (PDA, Shanghai Bio-way technology, Shanghai, China) and Vogel’s minimal media at room temperature under 12 h light/12 h dark conditions (21). Cultures were incubated for approximately 7 days, after which abundant orange conidia were produced and harvested for subsequent experiments.

Images of the hH1-sGFP strain were acquired using a 20× objective (NA 0.5) on a ZEISS AXIO Imager M2 microscope. GFP fluorescence was detected using excitation/emission (Ex/Em) wavelengths of approximately 488/509 nm. Scale bars are shown in each image and specified in the figure legends. Brightness adjustments were applied to the images using Zeiss ZEN microscopy software to optimize signal-to-noise and visualize weak signals without altering the original data.

### Chromosome counting

Conidia of *N. crassa* grown on PDA plates for 10 days were harvested and resuspended in 1.7 mL of protoplast buffer (0.8 M MgSO_4_, 0.2 M sodium citrate, pH 5.5) at a concentration of 10^7^ spores/mL. Nocodazole was added to a final concentration of 3 μg/mL to arrest the cell cycle. The suspension was incubated with shaking at 28 °C and 200 rpm for 1 hour. Cell wall digestion was then performed by adding 0.3 mL of an enzyme solution (1 M sorbitol, 50 mM sodium citrate, pH 5.8) containing 60 mg of Novozymes VinoTaste Pro, followed by incubation at 28 °C with shaking at 150 rpm for 1 hour.

The resulting protoplasts were pelleted by centrifugation at 5000 *g* for 10 minutes, washed once with 2 mL of 0.6 M KCl, and centrifuged again under the same conditions. The pellet was resuspended in 100 μL of 0.6 M KCl and fixed by adding 400 μL of a methanol–acetic acid solution (3:1, v/v). Fixed protoplasts were applied onto glass slides from a height of 10–15 cm and air-dried at room temperature.

Slides were stained with DAPI (1 μg/mL) and observed under a ZEISS Axioscope 5 fluorescence microscope with a 100× oil-immersion objective (NA 1.4) for chromosome counting. DAPI fluorescence was detected using Ex/Em wavelengths of approximately 353/465 nm. Upon slide preparation, microscopic examination indicated that fewer than 1% of the protoplasts ruptured, while the majority remained intact with clearly visible nuclei. Due to clustering of some protoplasts, which complicated the assignment of scattered chromosomes, only chromosomes unambiguously contained within a single, ruptured protoplast were included in the analysis. Chromosome numbers were recorded from a total of 105 such ruptured protoplasts and used to construct the chromosome distribution histogram.

### Monitoring chromosome division in live cells with confocal fluorescence microscopy

Conidia of *N. crassa* (hH1-sGFP strain) were harvested from PDA plates after 7–10 days at room temperature and resuspended in Vogel’s medium at a concentration of 10^6^ spores/mL. Suspensions were incubated for 3 h at 30℃ (225 rpm) to induce germination when active nuclear division begins to occur. Germlings were transferred to 384-well glass-bottom plates (Cellvis) for imaging.

Time-lapse imaging was performed on a Leica Stellaris 5 laser-scanning confocal microscope using a 63×/1.40 NA oil-immersion objective. Fields containing cells with 1–5 nuclei were selected (to avoid post-duplication hyphae). GFP was excited at 488 nm and emission collected at 495–530 nm with Tau gating (1–5 ns). Single-plane time series were acquired every 5 or 10 s at 200 Hz scan speed with five-line averaging and a frame size of 512×512 or 1024×512 pixels. Chromosome counts were estimated from condensed chromosome morphology at metaphase/early anaphase. Because chromosomes reorient rapidly during division, counts were refined by examining multiple time points in the series, during which the event presented different orientations; the focal plane was manually adjusted as needed to maintain optimal focus on chromosomes. Image processing and analyses were performed with Fiji (ImageJ, v1.54p).

### Relative fluorescence intensity (RFI) determination of nuclear DNA content by flow cytometry

To measure the DNA content within nuclei, flow cytometry was performed following a previous study with minor modifications, using nuclei isolated from stationary-phase *Saccharomyces cerevisiae* liquid cultures and from mycelia of *S. sclerotiorum* and *N. crassa* (1). Briefly, fresh mycelia were collected and grounded into fine powder with liquid nitrogen. Approximately 50 mg of the resulting material was resuspended in 2 mL of homogenization buffer [10 mM PIPES (Catalog #AAA1609014; Fisher Scientific), 5 mM CaCl2, 5 mM MgSO4 and 0.5 M sucrose]. Samples were centrifuged at 8,000 *g* for 20 minutes at 4℃ to pellet nuclei. The nuclei-containing pelleted was resuspended in homogenization buffer and centrifuged at 100 *g* for 5 minutes. Next, 1.8 mL of supernatant was transferred to a new tube and centrifuged again at 8,000 *g* for 20 minutes. The final pellet, enriched for nuclei, was resuspended in 2 mL of cold sterile 1× PBS buffer (Catalog # B548117; Sangon Biotech).

For fluorescence activated cell sorter (FACS) analysis, isolated nuclei were stained with 50 μg/mL propidium iodide (PI) for 5-10 minutes. FACS was performed using a Sony MA900 Multi-Application cell sorter instrument (Sony Biotechnology, Japan) equipped with a 100 μm sorting chip (LE-C3210), a 488 nm excitation laser, and a 617/30 nm emission detector. Nuclei were gated based on BSC-A (backscatter area) and PI-A (PI fluorescence area) to distinguish populations by size and fluorescence intensity. The sorting was validated by examining sorted nuclei under a fluorescence microscope. Nuclei isolated from S. *cerevisiae, S. sclerotiorum* and *N. crassa* mycelia were sorted into 96-well plates, with each species occupying 15 wells and each well containing 100 nuclei. Mean fluorescence intensity values from each well were recorded and used for comparative analysis across samples.

### FISH analyses

Through BLAST searches of the *N. crassa* genome in FungiDB and EnsemblFungi, a linkage group I (LG I)-specific probe was identified (5’ Cy5-CTAGCACCCTTCACTAGCACCCTTCACTAGCACCCTTCA 3’). This specific probe is located at around 7.52 Mb in a non-coding region with a single successive hybridization site, which lies outside of the defined heterochromatin block. In addition, a linkage group IV (LG IV)-specific probe was identified from literature (22), which targets a region around 2.6–2.7 Mb and is not located near sub-telomeric heterochromatic regions (5’ Cy5-GTGAACCATCGAGT**Y**TTTGAACGCAAGT 3’). Additionally, a telomere repeat probe capable of targeting all chromosomes and containing multiple adjacent binding sites was identified (5’ Cy5-CCCTAACCCTAACCCTAACCCTAACCCTAA 3’). Probes with a 5’ Cy5 modification were synthesized by Integrated DNA Technologies (IDT).

Conidia fixation and FISH were performed following our previous study with minor changes (1). Briefly, the conidia were fixed in an ethanol-acetic acid solution (3:1, v/v) for 3 hours, followed by dehydration through a graded cold ethanol series of 70%, 90%, and 100%, with 5-minute intervals. The samples were then treated with RNase (100 μg/mL in 2× SSC) in a humid chamber at 37℃ for 1 hour and subsequently washed four times with 2× SSC at room temperature, with 2-minute intervals. FISH hybridization solution (50% formamide, 10% dextran sulfate, and 2× SSC) was mixed with 100 ng of probe. After adding the hybridization mix, the slides were sealed with a glue gun to prevent evaporation, wrapped in aluminum foil and incubated at 80℃ for 10 minutes to denature genomic DNA. They were then transferred to a moist, dark chamber for hybridization. Hybridization with the nonspecific probe was performed overnight at 37–42 °C, whereas hybridization with the LG I-specific and LG IV-specific probes was carried out for 22 hours at 45 °C and 35–40 °C, respectively. The slides were sequentially washed with 2× SSC at room temperature for 5 minutes, 0.1× SSC at 42–45 ℃ for 10 minutes, and 2× SSC at room temperature 5 minutes, followed by dehydration through the cold ethanol series. Prior to imaging, 20 μl of 1 μg/mL DAPI (Sigma-Aldrich, Canada) was added to stain nuclei. 5 μl of VECTASHIELD HardSet Antifade Mounting Medium (Vector Labs, USA) was applied, and the slide was sealed with clear nail polish (Sally Hansen 109 invisible, Canada) to preserve fluorescence. The cells without signals inside the nuclei are considered samples failed the experiment (column “Number of conidia without signals inside the nuclei”).

Images of FISH samples were acquired using a 100× oil-immersion objective (NA 1.4) on a ZEISS AXIO Imager M2 microscope. DAPI (Ex/Em ≈ 353/465) and Cy5 (Ex/Em ≈ 650/673) fluorescence were used. Scale bars are shown in each image and specified in the figure legends. Brightness adjustments were applied to the images using Zeiss ZEN microscopy software to optimize signal-to-noise and visualize weak signals without altering the original data.

## Acknowledgments

We thank Josh Li for assistance *N. crassa* conidia cultivation, and Dr. Lei Tian and Xuanye Chen for careful reading of the manuscript. We are grateful to Prof. Yi Yang (SU) for sharing their ZEISS fluorescence microscope for chromosome counting. ChatGPT 5.0 (OpenAI, San Francisco, CA, USA) was used for correcting grammar mistakes and language polishing. This work was supported by funds to Y.X. from National Key Research and Development Program of China (2025YFF1000600) and to X.L. from the Canadian Natural Sciences and Engineering Research Council (NSERC) Discovery program, NSERC-CREATE-PRoTECT, Canada Research Chair (CRC), and the Canadian Foundation for Innovation (CFI) funds. J.T. is partly supported by a UBC Four-year Fellowship.

## Supporting information

**S1 Movie. Live chromosome division in *N. crassa* conidia**.

Germinating conidia expressing hH1–sGFP were imaged by laser-scanning confocal microscopy at room temperature. Frames highlight anaphase segregation with an estimated two chromosomes visible. Scale bar, 2 µm. The GFP channel is shown in (A), and the merged channel is shown in (B).

**S2 Movie. Live chromosome division in *N. crassa* young mycelia**.

A young mycelium tip germinated from a conidium expressing hH1–sGFP was imaged by laser-scanning confocal microscopy at room temperature. Frames highlight prophase condensation, with an estimated number of four chromosomes visible. Scale bar, 2 µm. The GFP channel is shown in (A), and the merged channel is shown in (B).

## Notes

### Competing Interest Statement

The authors have declared no competing interest.

### Summary of Updates

Additional experiments were conducted, including chromosome counting, live-cell imaging, and supplementary FISH analyses, and the conclusions were revised accordingly.

